# Metabolic organization of macaque visual cortex reflects retinotopic eccentricity and category selectivity

**DOI:** 10.1101/2025.09.27.678945

**Authors:** Hiroki Oishi, Vladimir K. Berezovskii, Margaret S. Livingstone, Kevin S. Weiner, Michael J. Arcaro

## Abstract

Neural activity requires energy metabolism, and the brain’s metabolic architecture varies across regions. Yet, it remains unclear whether these variations are meaningfully related to the functional and perceptual demands of cortical processing. In high-level visual cortex, category-selective regions, such as those preferentially responding to faces and scenes, are consistently distributed along a topographic axis that varies in sensitivity to spatial scale, a feature dimension that imposes differing metabolic demands at the level of the retina. This axis reflects a broader organizing principle of the visual system: retinotopic eccentricity, the topographic mapping of visual space relative to gaze. Here, we tested whether cortical metabolic architecture reflects this principle by aligning *in vivo* fMRI maps of eccentricity and visual category selectivity with *ex vivo* cytochrome oxidase (CO) histology, a marker of oxidative metabolism, in macaque visual cortex. We found that face-selective region ML, which is biased toward central vision, exhibits higher CO intensity than the peripherally-biased scene-selective region LPP. More broadly, CO intensity covaries with eccentricity across the entire occipitotemporal cortex, though this gradient only partially accounts for the elevated CO in ML. These findings reveal a close correspondence between cortical energy consumption and retinotopic representation, suggesting that metabolic resources are shaped by the processing demands of visual perception.

## Introduction

Neural activity underlying perception, cognition, and behavior is sustained by cortical energy metabolism. Despite the brain’s high energetic demands (1), the principles governing how metabolic resources are distributed across cortical systems remain a central challenge in neuroscience. Recent studies in both humans and nonhuman primates have revealed that the cortical metabolic architecture is not spatially uniform on a large scale (2, 3). However, it remains unclear whether these variations are meaningfully related to the brain’s functional architecture, particularly the large-scale systems that support behaviorally relevant computations. We hypothesize that cortical energy metabolism is shaped by the specific processing demands of these functional systems.

Cytochrome oxidase (CO) histochemistry has long served as a key method for mapping cortical metabolic architecture, revealing discrete functional compartments across sensory systems (4). Variations in CO intensity reflect differences in metabolic demand that have been linked to neural activity in early sensory cortices, such as CO-rich “blobs” in primary visual area V1 (5, 6), barrels in somatosensory cortex (7), and tonotopically organized modules in auditory cortex (8). These modules often coincide with high spontaneous firing rates and high metabolic demand of local circuits that are specialized for high-acuity or precision-tuned processing. Prominent staining in intermediate layers of primary cortices, such as layer IVc of V1 or layer IV of S1, also reflects the high metabolic costs of dense thalamic input processing. While this body of work has established the utility of CO staining for revealing functional subdivisions at the mesoscale, particularly in early sensory areas, much less is known about how metabolic architecture is distributed across cortical regions and hierarchies. In particular, the relationship between early metabolic specialization and downstream stages of processing, and whether large-scale functional systems exhibit corresponding patterns of energy metabolism, remains largely unexplored.

Inferior temporal (IT) cortex offers a strong test case for addressing this question. IT supports high-level visual recognition and contains spatially organized domains that vary in the spatial scale and acuity of the input they process, reflecting tuning to different perceptual demands. A well-studied example is the contrast between face- and scene-selective regions. Face-selective areas support fine-grained analysis of local features within central vision, critical for tasks like identity recognition and expression decoding. In contrast, scene-selective regions are more sensitive to the global configuration and spatial layout of the environment, integrating information across the visual periphery to support navigation and context perception (9–11). These areas are not randomly distributed but instead align along a continuous topographic axis of retinotopic eccentricity, the mapping of visual space relative to gaze, that extends from the early visual cortex through IT. This axis provides a consistent large-scale scaffold for functional specialization in the visual system and presents a principled framework for testing whether metabolic architecture reflects retinotopic organization.

Here, we investigated whether cortical metabolic architecture reflects large-scale functional organization in the primate visual system, focusing on its relationship to both retinotopic eccentricity and visual category selectivity. We combined *in vivo* fMRI mapping of eccentricity and category selectivity with *ex vivo* CO histology in the macaque visual cortex. We first compared CO intensity between two functionally and retinotopically distinct regions in IT: the face-selective patch ML, which emphasizes central visual input, and the periphery-biased scene-selective patch LPP. Across the ventral visual pathway and within each region, CO intensity covaried with eccentricity representation in superficial and deep segments of cortex, revealing a continuous metabolic gradient aligned with the system’s topographic organization. Notably, ML exhibited elevated CO levels not only relative to LPP but also compared to other visual areas, suggesting region-specific metabolic demands beyond eccentricity. These findings demonstrate that cortical metabolic architecture reflects functional layout, shaped by both the spatial structure of visual input and the processing demands of distinct visual domains.

## Results

### CO architecture differs between face- and scene-selective regions

To assess metabolic architecture in the visual cortex, we leveraged a multimodal dataset combining *in vivo* fMRI and *ex vivo* histology from two macaque monkeys. This dataset was previously used to examine variation in cytochrome oxidase (CO) intensity across fMRI-defined face-selective patches in inferior temporal (IT) cortex, focusing on comparisons within a functionally coherent network (2). In the present study, we extended this approach to test whether CO variation also reflects differences across functionally distinct networks within IT. Specifically, we compared CO intensity between two fMRI-defined regions with divergent category selectivity (Figure 1A). One region, the middle lateral face patch (ML), was defined by strong neural responses to images of faces and weak responses to images of objects and scenes. The other region, the lateral place patch (LPP), responded strongly to scenes and weakly to images of objects and faces. Among the face-selective regions, we focused on ML because it is located at a similar anterior–posterior position as LPP, enabling a direct comparison while minimizing potential bias from the CO gradient that varies along this axis of IT cortex (2). Comparisons involving more anterior face patches would be confounded by this gradient, making it difficult to attribute differences in CO intensity to category selectivity rather than anatomical position.

**Fig. 1.**
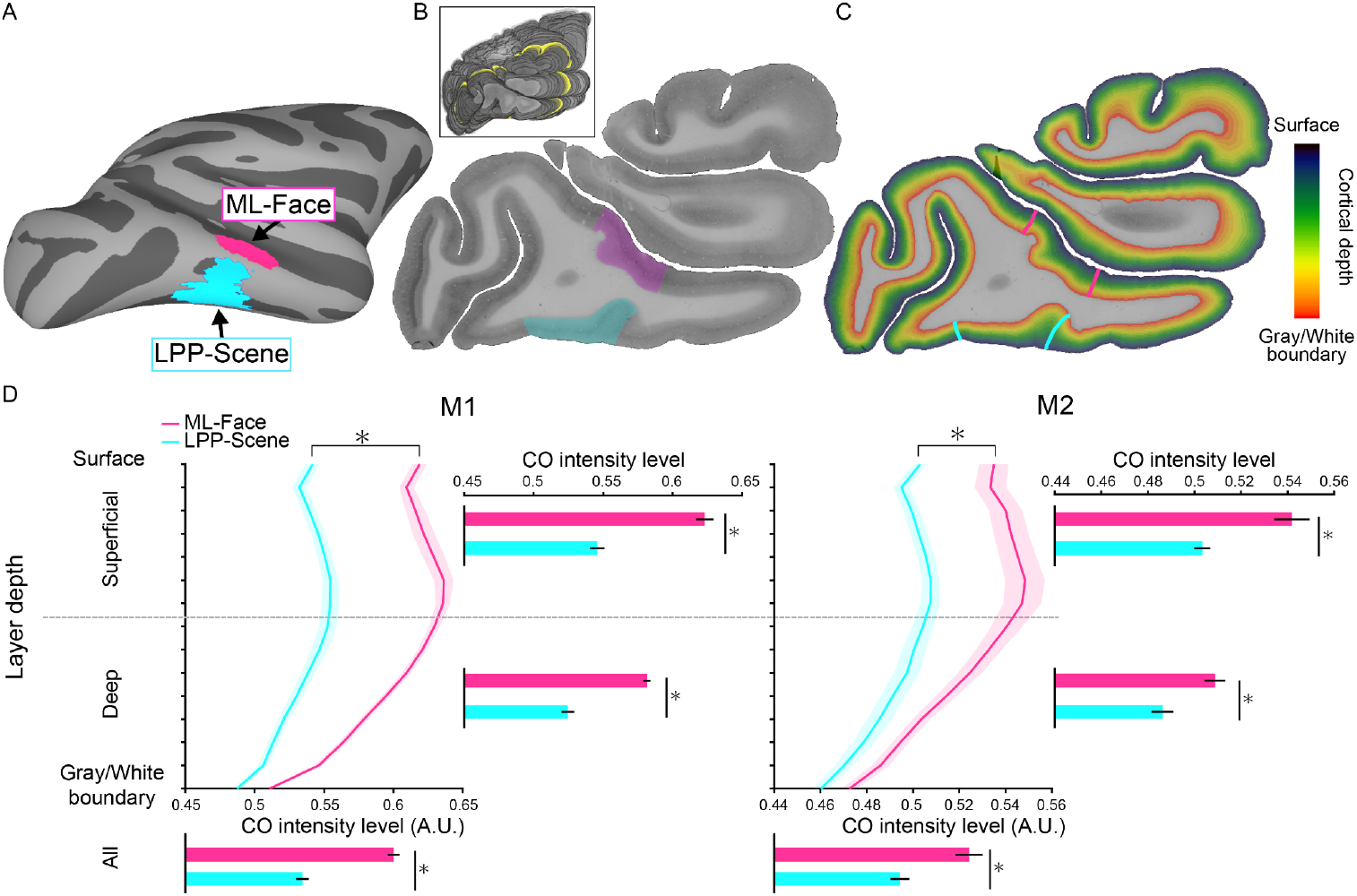
Cytochrome oxidase (CO) architecture differs between face and place patches. (A). A cortical surface of the right hemisphere, illustrating the anatomical locations of face-selective patch ML (magenta) and scene-selective patch LPP (cyan) from probabilistic maps (n = 6 for ML, n = 9 for LPP). (B). The ML and LPP localized in a representative histological section stained for CO. The ML and LPP are colored in translucent magenta and cyan, respectively. The inset image shows the 3D histological volume created by coregistering serial sections. The yellow-highlighted section corresponds to the representative section. The ML and LPP were projected onto histological sections by aligning the 3D anatomical MRI to the histological volume. (C). Cortical depth maps. Colored lines illustrate the boundaries of ML and LPP. (D). Cortical depth profiles of CO intensity are shown for ML and LPP in both hemispheres of monkey M1 (left) and M2 (right). Curves represent average CO intensity, with shading indicating +/-1 SEM across histological sections. Bar plots to the right of each depth profile show mean CO intensity for superficial (top seven bins) and deep (bottom eight bins) portions of cortex. Mean CO intensity across the full cortical depth is shown below each profile. Asterisks indicate significant differences between ML and LPP (p < 0.01, FDR-corrected). Higher CO values indicate darker staining.

As described previously (2), cortical histological sections were reconstructed into 3D volumes by stacking and aligning adjacent 50-μm sagittal sections stained for CO (Figure 1B). Functional regions of interest (ROIs) were defined using probabilistic maps of category selectivity from fMRI and aligned to the histological volume through intermediate registration to a high-resolution anatomical MRI (See Methods). To characterize the distribution of CO across cortical depth, each section was segmented into 15 equal-volume bins at varying depths parallel to the cortical surface (Figure 1C). While these bins do not correspond to cytoarchitectonic layers, they span the full thickness of the gray matter, allowing depth-resolved quantification of staining intensity.

To test whether category-selective regions differ in metabolic architecture, we compared the CO depth profiles between ML and LPP. In both regions, CO intensity increased from superficial to middle depths and decreased toward deeper portions of cortex (Figure 1D), with peak staining observed in relatively superficial portions. While the overall CO depth profile was similar between ML and LPP, ML exhibited significantly higher CO intensity than LPP when averaged across the full cortical depth, as well as within both superficial and deep portions (all ts > 3.104, all ps < 0.004, FDR-corrected; Figure 1D). These findings demonstrate that face- and scene-selective regions in IT differ in metabolic staining intensity, and that this difference spans the entire cortical depth.

### CO intensity correlates with eccentricity in both face- and scene-selective regions

The observed differences in CO intensity between ML and LPP may reflect a categorical distinction between face and scene processing. However, an alternative or complementary explanation is that they reflect differences in retinotopic eccentricity, which varies both within and between these regions. Spatial resolution varies systematically with eccentricity, with central vision supporting finer discriminations than the periphery. For example, face perception relies on high-acuity central vision, while scene perception draws on coarser information across the visual field. In the retina, cone-dense foveal regions supporting fine spatial vision show higher metabolic demands than rod-dominated peripheral regions. If this relationship carries forward to the visual cortex, including downstream IT cortex, it raises the possibility that CO intensity may covary with eccentricity representation within each region.

To test this, we projected probabilistic maps of retinotopic eccentricity from fMRI onto the histological sections (Figure 2A). Within each ROI, the cortex was divided into spatial bins oriented orthogonal to the cortical surface. Although these bins do not correspond to anatomical cortical columns, they allowed localized sampling along the cortical ribbon. For each bin, we averaged CO intensity across cortical depth and extracted the corresponding eccentricity value (Figure 2B and 2C). This enabled direct comparison between local metabolic activity and retinotopic organization. Within both ML and LPP, CO intensity was significantly correlated with eccentricity (Pearson’s r = −0.475 and −0.639 for ML and LPP, respectively; Figure 2D), with higher CO intensities in portions of cortex representing central visual space. These results indicate that differences in retinotopic organization contribute to the metabolic variation observed even within downstream, category-selective cortex at the apex of the visual processing hierarchy.

**Fig. 2.**
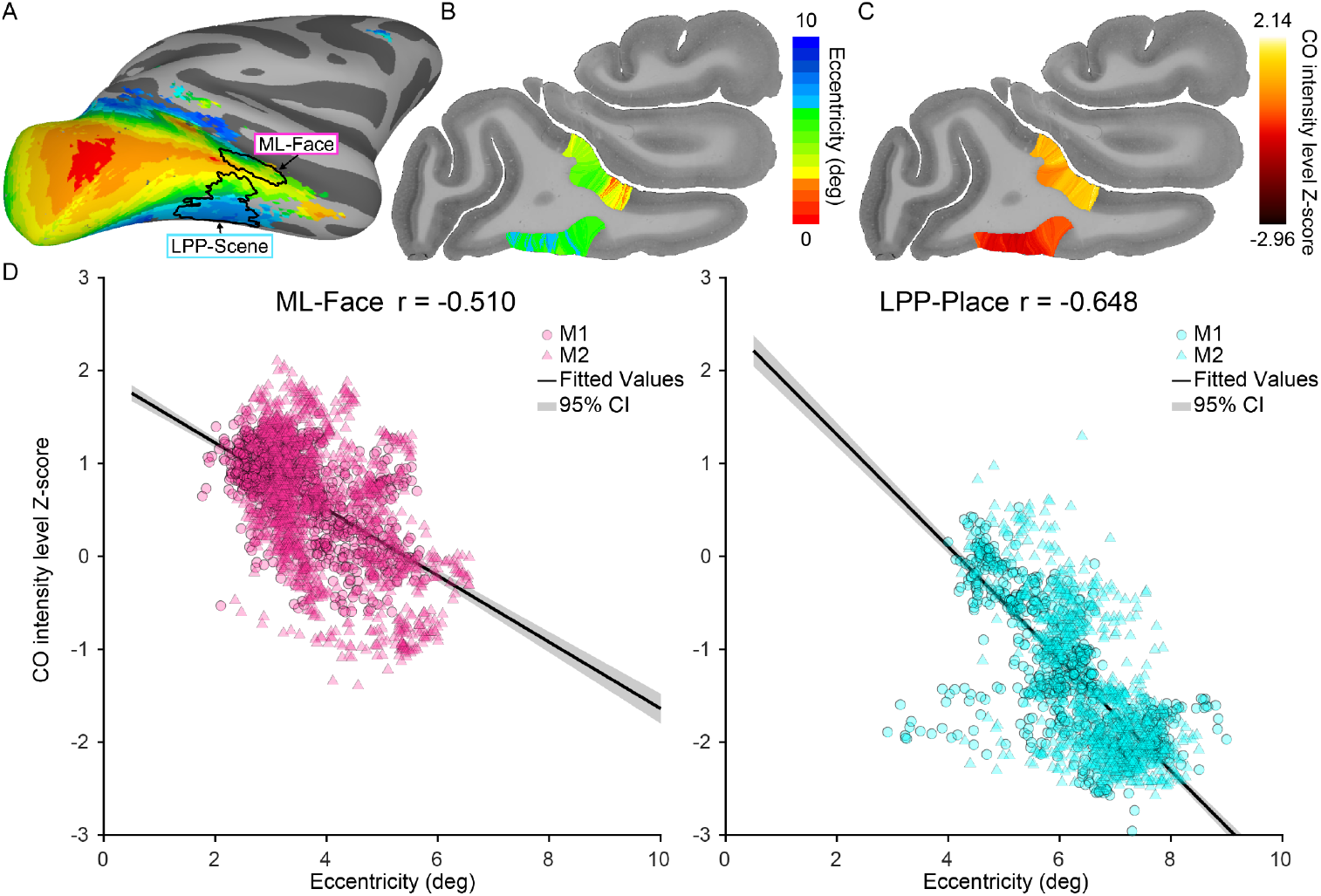
CO intensities are correlated with eccentricity in face and place patches. (A). A group average eccentricity map (n = 6) shown in the same surface view as Figure 1A. The color code indicates the preferred eccentricity at each cortical location, covering the central 10 degrees of visual space from center (red/yellow) to periphery (blue). The outlines of ML and LPP are overlaid on the cortical surface. (B). A representative histological section, illustrating the eccentricity representations in ML and LPP. Within ML and LPP, the cortex was divided into spatial bins oriented orthogonally to the cortical surface, and an eccentricity value was assigned to each bin based on the eccentricity map projected onto the histological data. (C). Mean CO intensities within the spatial bins in ML and LPP. CO intensities across cortical depth were averaged for each bin. (D). CO is correlated with eccentricity in ML (left) and LPP (right). Each point represents a spatial bin within the ROI (ML, magenta; LPP, cyan). Data from M1 and M2 were Z-scored separately (M1: circles; M2: squares). Black lines show linear regression fits with gray shading indicating 95% confidence intervals.

### Eccentricity predicts CO intensity across visual cortex

Having established a local relationship between CO intensity and eccentricity within both face- and scene-selective regions, we next asked whether this relationship extends more broadly across the visual cortex. To address this, we expanded the spatial sampling analysis, dividing the full extent of the occipitotemporal cortex into bins oriented orthogonal to the cortical surface. For each bin, we derived average CO intensity and eccentricity values across cortical depth. This analysis revealed a significant negative correlation between CO and eccentricity in both monkeys (Pearson’s r = −0.572 for M1 and −0.441 for M2; Figure 3), indicating that regions representing central visual space consistently exhibit higher metabolic activity. These findings suggest that the relationship between CO intensity and eccentricity is a fundamental organizational feature across the visual hierarchy.

**Fig. 3.**
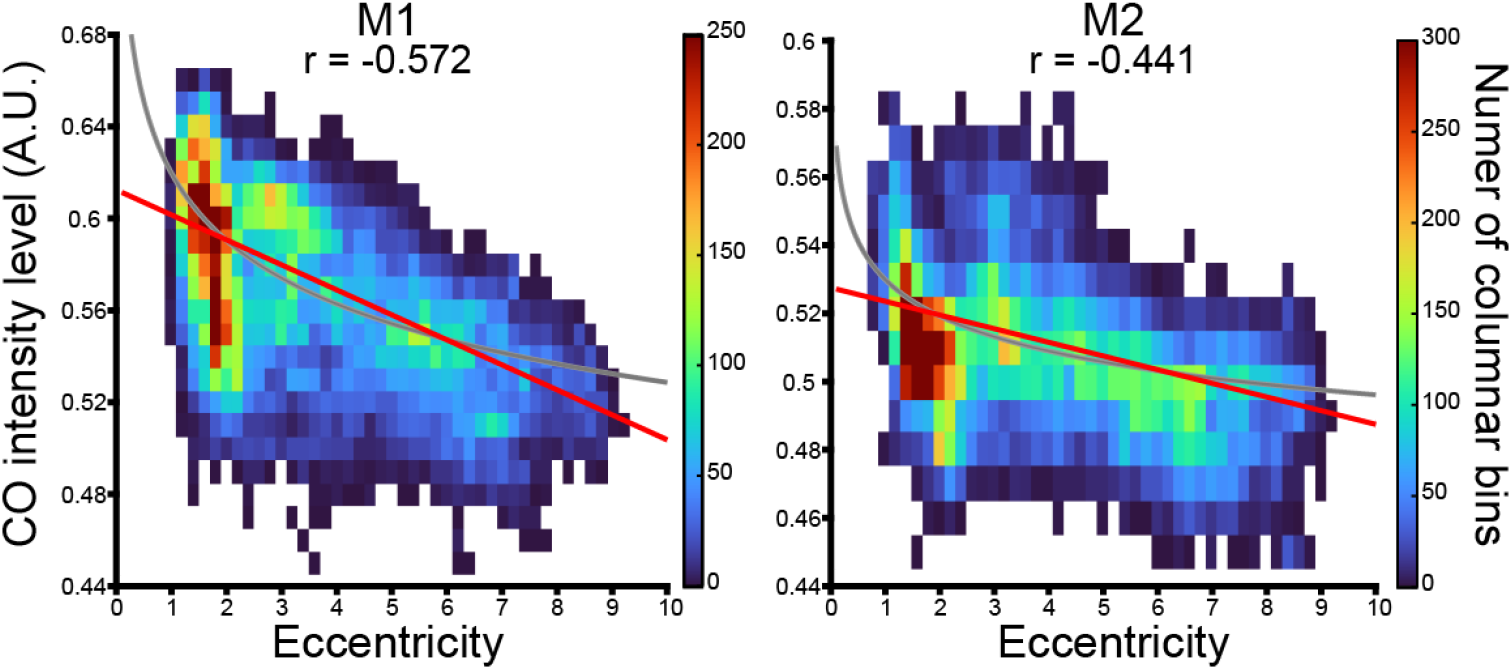
CO intensity varies systematically with eccentricity across the occipitotemporal visual cortex. Two-dimensional histograms show the distribution of spatial bins across occipitotemporal cortex as a function of eccentricity (bin width = 0.2 deg) and a CO intensity (bin width = 0.01) for monkey M1 (left) and M2 (right). Warmer colors indicate a higher number of spatial bins within each bin pair. Red lines indicate linear regression fits, and gray curves show low-order polynomial fits. In both monkeys, CO intensity is significantly correlated with eccentricity across the visual cortex.

To examine how this CO-eccentricity relationship varies across the ventral visual hierarchy, we projected the probabilistic atlas of occipitotemporal retinotopic areas from V1 to PITd/v onto the histological sections. For each area, we then calculated the mean CO intensity and the mean eccentricity value to assess how these metrics change across the visual hierarchy. Two patterns emerged: This posterior-to-anterior decrease in CO was accompanied by a shift toward more peripheral eccentricity values, consistent with the known reduction in cortical magnification from V1 to higher visual areas (12, 13). These relationships were apparent for both superficial and deeper portions of cortex (Pearson’s r = −0.406 for superficial and −0.519 for deep; Figure 4A), indicating that the CO-eccentricity relationship spans the entire cortical depth. Together, these findings reveal a systematic variation in metabolic activity across the visual hierarchy that is closely tied to retinotopic eccentricity.

**Fig. 4.**
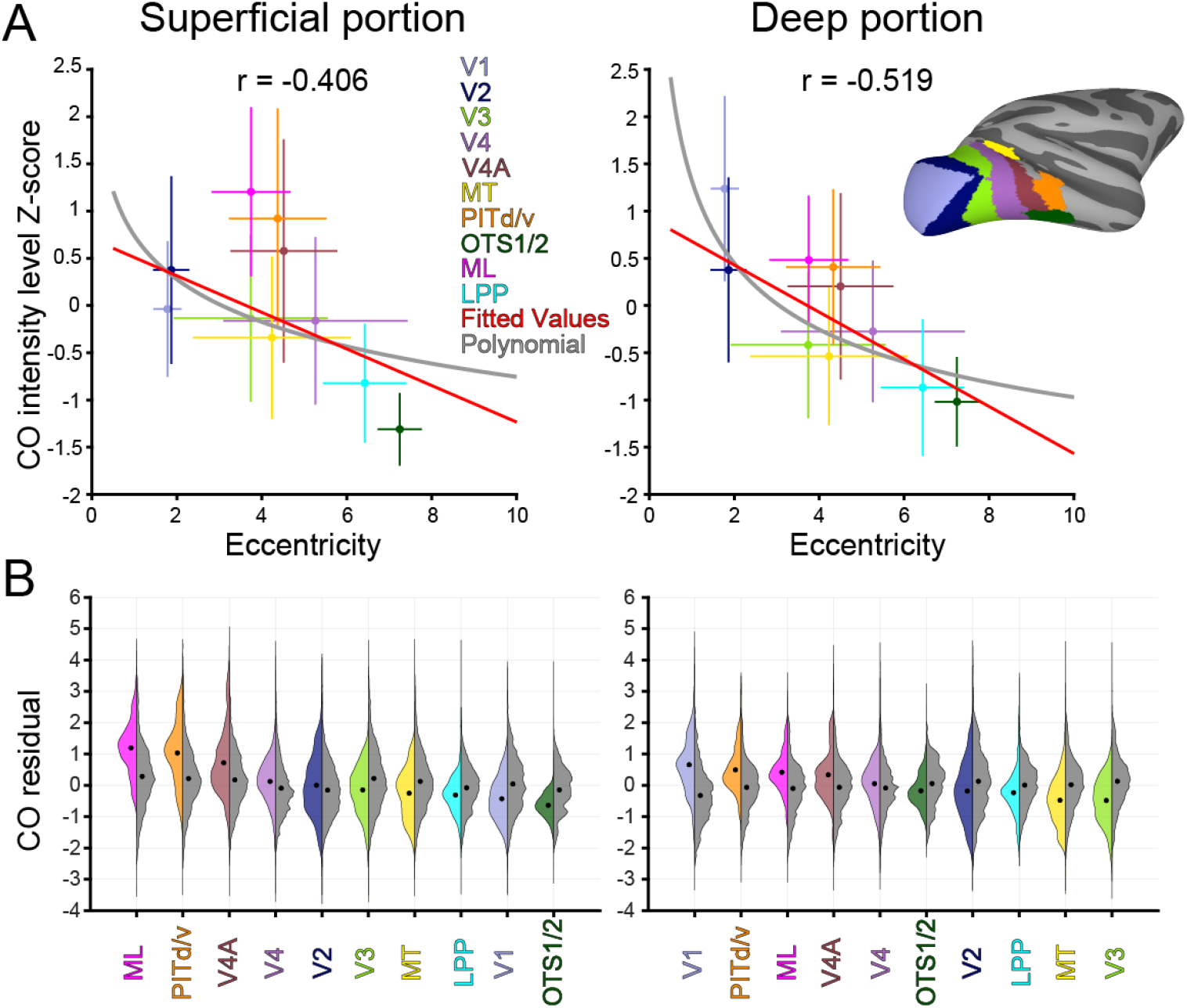
CO intensity variation across occipitotemporal areas reflects eccentricity biases, with additional effects in face-selective regions and input layers of V1. (A). Mean CO intensity and eccentricity for retinotopically defined visual areas are shown separately for superficial (left) and deep (right) portions of cortex. Each point represents the average value for one visual area, with error bars indicating standard deviation across spatial bins. Areas include V1, V2, V3, V4, V4A, MT, PITd/v, and OTS1/2. The colors of the points match the area labels shown in the left panel. The cortical surface in the right panel displays the anatomical locations of the visual areas. For reference, the points of ML (magenta) and LPP (cyan) are also shown; ML largely overlaps with PITd/v, and LPP with OTS1/2. Linear (red) and polynomial (gray) fits summarize the relationship between mean eccentricity and mean CO intensity across areas. (B). CO residual distributions for visual areas in superficial (left) and deep portions. Split violin plots comparing CO residual distributions for each visual area (left, colored by ROI) against eccentricity-matched null distributions (right, gray). Residuals represent deviations from the global robust linear regression of CO intensity on eccentricity. Null distributions were created by sampling voxels from all other visual areas with identical eccentricity profiles (5000 iterations). Areas are ordered by mean residual magnitude (highest to lowest). Black circles indicate mean values for each distribution.

### Elevated CO in ML is not explained by central visual field bias

Although ML and LPP differ in category selectivity, they also differ in retinotopic bias, with ML representing more central and LPP more peripheral visual space. Because regions that represent central vision tend to show elevated CO intensity across the visual cortex, it is possible that the higher CO levels observed in ML simply reflect this central visual field bias. To test this, we used the linear model relating CO intensity to eccentricity described above and computed the standardized residuals for each columnar bin. We assessed whether individual ROIs deviated meaningfully from the global relationship between CO and eccentricity. We applied an eccentricity-matched null approach, which tests whether each area’s residual distribution differs meaningfully from what would be expected based on sampling from the broader visual cortex at matched eccentricities. For each ROI, we constructed a null distribution by sampling residual values from all other visual areas at precisely matched eccentricity distributions, repeated 5000 times to ensure stability (see Materials and Methods).

This analysis revealed striking depth-specific patterns of metabolic organization (Figure 4B, Fig. S1 and Table S1). In deep cortical depth portions, V1 showed the most pronounced deviation from eccentricity expectations, with an adjusted effect size of 1.07 (95% CI: 1.035 - 1.105), explaining 23.3% of variance and showing only 54.5% distributional overlap with its matched null. This pattern is consistent with the high metabolic demands of Layer IVC, which receives dense LGN input. ML also showed elevated CO in deep portions (adjusted d = 0.816, 12.5% variance explained) but with greater distributional overlap (66.4%). In contrast, LPP showed a modest negative deviation in deep portions (adjusted d = −0.376) with high overlap (74.8%) and minimal variance explained (3.3%), indicating its residual distribution was largely indistinguishable from the eccentricity-matched null. In superficial portions, the pattern shifted dramatically. ML emerged as the strongest outlier (adjusted d = 1.10, 19.1% variance explained, 57.6% overlap), while V1 showed a negative deviation (adjusted d = −0.698, 8.0% variance explained). LPP again showed minimal meaningful deviation (adjusted d = −0.452, 3.4% variance explained, 79.3% overlap), suggesting its CO patterns largely conform to retinotopic expectations. These findings demonstrate that the elevated CO in ML cannot be attributed solely to its central visual field bias, as it shows pronounced deviations even when compared to other areas with matched eccentricity distributions.

### Category and eccentricity effects are not explained by tissue size

Although our analyses revealed strong relationships between CO intensity, category selectivity, and eccentricity, these effects could be influenced by systematic variability in staining across histological sections. Such variation can arise from technical artifacts during processing. Differences in tissue size across sections can lead to uneven staining, since smaller sections may absorb more stain when incubated for a fixed amount of time. Indeed, we observed a strong correlation between tissue size and CO intensity of the sections in both animals (Pearson’s r = −0.690 for M1 and −0.775 for M2). To control for this potential confound, we regressed CO intensity against tissue size across all columnar bins within the occipitotemporal cortex. We then used the residuals from this model, which reflect variation in CO orthogonal to tissue size, for all subsequent analyses. The differences in depth profiles between ML and LPP remained significant (all ts > 2.721, all ps < 0.010, FDR-corrected), as well as the correlations between CO intensity and eccentricity in ML and LPP (Pearson’s r = −0.672 and −0.617 for ML and LPP, respectively). The broader relationship between CO intensity and eccentricity across the visual cortex was also preserved (Pearson’s r = −0.342 for M1 and −0.303 for M2; r = −0.312 for superficial and −0.406 for deep). These findings confirm that the observed CO gradients and their functional associations are not artifacts of staining variability, but instead reflect intrinsic cortical organization.

## Discussion

Cortical specializations differ in both anatomical architecture and energetic demand. These differences are not randomly distributed, but reflect underlying organizational principles that structure the brain’s layout. In the visual system, one such principle is retinotopic eccentricity, which organizes cortical representations according to distance from the center of gaze and covaries with spatial resolution, receptive field properties, and category selectivity. Here, we show that cortical metabolic architecture, as indexed by cytochrome oxidase (CO) intensity, also aligns systematically with eccentricity. Face- and scene-selective regions in inferotemporal cortex (IT) differ in CO intensity, and within each region, CO values covary with eccentricity. More broadly, a continuous CO–eccentricity gradient spans the occipitotemporal visual cortex. These results suggest that energy use in the cortex is distributed according to the spatial structure of the visual input and the processing demands it imposes.

Why might regions representing central vision be more metabolically demanding? In the retina, cones dominate the fovea and support high-acuity vision, but they do so at a steep energetic cost (14–16). Their higher mitochondrial content, sustained ion exchange, and increased synaptic transmission result in elevated oxidative metabolism relative to peripheral rods (17, 18). This creates a structured metabolic gradient from the outset of visual processing. Our findings suggest that this gradient is preserved across the visual hierarchy, with central visual field representations consistently showing higher CO intensity throughout the cortex. Although the cellular architecture differs between retina and cortex, both appear to follow a shared principle in which fine-grained visual processing carries consistent metabolic costs.

This raises the question of whether eccentricity-dependent metabolic architecture plays a formative role in cortical specialization or whether it simply reflects the demands of ongoing visual behavior. In macaques, adult-like CO patterns in V1 are present at birth and have been linked to emerging functional distinctions (19) and have been linked to emerging functional distinctions (e.g., (20)). While the organization of CO in the higher visual cortex at birth remains unstudied, retinotopic maps are well established before the appearance of category-selective regions in IT (10). This developmental sequence suggests that early spatial and metabolic biases could constrain where functional specializations emerge. At the same time, visual experience likely contributes to metabolic organization. For example, infants spend a disproportionate amount of their early looking time fixating on faces, concentrating visual stimulation within central vision and potentially reinforcing eccentricity-related metabolic gradients (21). This interpretation is supported by modeling work in which eccentricity-dependent energetic constraints gave rise to increased energy use in central regions, mirroring the cortical gradient observed in our data (22). These perspectives are not mutually exclusive. Early metabolic biases may help guide the formation of specialized regions and remain embedded in the adult cortex, where they continue to support the demands of mature vision.

Although our findings reveal a strong relationship between eccentricity and CO intensity, this relationship may be shaped by multiple co-aligned factors. Eccentricity covaries with several functionally relevant properties, including receptive field size, spatial frequency tuning, curvature sensitivity, and category selectivity (23–26). As a result, CO gradients may reflect the cumulative metabolic cost of fine-scale processing rather than eccentricity alone. For a given fixation, neurons representing central visual space must resolve denser contrasts and finer details than those representing peripheral space, which imposes a greater energetic burden. This provides a straightforward functional explanation for why regions encoding central vision show elevated CO intensity. Moreover, CO intensity is not necessarily driven solely by local cortical activity. In V1, CO-rich blobs correspond to the terminal zones of koniocellular inputs from the LGN, and similar projections from the pulvinar may underlie extrastriate CO compartments (27, 28). These observations raise the possibility that thalamocortical input contributes to metabolic architecture in ways that are independent of intrinsic cortical processing. Future work should further disentangle these factors. Studies of individuals with atypical visual experience, such as developmental prosopagnosia, or experimental manipulations in animals, such as deprivation of specific categories or training with novel stimuli (29), could clarify how category experience shapes metabolic architecture. Integrating CO histology with measures of thalamocortical connectivity and developmental time points will also help determine how metabolic gradients arise and are maintained. Our study focused on the ventral visual pathway, but the dorsal stream also contains well-characterized retinotopic eccentricity maps (30–32). Assessing whether the parietal cortex exhibits similar metabolic gradients would provide a critical comparison, revealing whether energy use is structured by common spatial principles across visual systems.

More broadly, our findings demonstrate the value of linking functional maps with metabolic architecture to uncover large-scale organizational principles of the brain. Neural computation is inherently metabolically expensive, yet few studies have directly examined how energy use is organized in relation to information-processing demands. By combining *in vivo* fMRI with *ex vivo* CO histology, we establish a direct link between behaviorally relevant cortical maps and their underlying metabolic architecture. Extending this approach to other sensory systems, as well as motor and cognitive domains, may reveal how structural and energetic constraints influence the development and organization of specialized brain functions. Ultimately, characterizing the energetic landscape of the brain may offer new insights into the biological cost of cognition and the pressures that shape its organization.

## Materials and Methods

### Non-human primates

All training, surgery, and experimental procedures were approved by the Harvard Medical School Animal Care and Use Committee and conformed with NIH guidelines for the humane care and use of laboratory animals. Two normally reared adult monkeys (Macaca mulatta) underwent T1w MRI and histological experiments. Nine normally reared adult monkeys underwent T1w and functional MRI.

### Histology and preprocessing

To quantify the neurometabolic architecture across the visual cortex, we analyzed the cytochrome oxidase (CO) staining data of the *ex vivo* two macaque monkeys, acquired in our previous study (2). As previously reported (33), monkeys that had reached endpoints were euthanized by intravenous injection of SomnaSol (120 mg/kg sodium pentobarbital), and transcardially perfused with a rinse solution (0.9% sodium chloride and 0.5% sodium nitrite) followed by 4% paraformaldehyde in 0.1 M phosphate buffer (pH 7.4). After overnight postfixation, brains were transferred into 30% sucrose in 0.1 M phenobarbital. Right hemisphere tissue was sectioned sagittally at 50-μm thickness using a freezing microtome, covering visual areas from V1 to IT. The left hemisphere was sectioned in a coronal orientation, only covering a small part of the IT anterior to V4, and was not included in this study. Serial sections were mounted on glass slides, postfixed in formol saline (10% formalin with 9 g/L sodium chloride) for 12 days, and processed for CO using standard protocols with catalase, cytochrome *c* and 3,3′-diaminobenzidine reagents.

Digital images of CO-stained sections were captured using a Panasonic Lumix DMC-ZS7 camera with a 12X optical zoom under uniform lighting with a light box. Images were converted to 8-bit grayscale and normalized from 0–255 to 0–1 range, where 0 indicates no CO staining and 1 indicates the darkest staining. To correct for variability in the staining intensity across slices, we used white matter as a within-slice reference. For each section, we subtracted the difference between the section’s mean white matter intensity and each gray matter pixel’s intensity. To preserve the original intensity scale, we then added back the grand mean white matter intensity averaged across all sections.

Cortical CO architectures were quantified as a function of location on the cortical surface and depth by segmenting the gray matter into small spatial units using LayNii (https://github.com/layerfMRI/LAYNII; 34). For each sagittal section, we used the LN_COLUMNAR_DIST tool to generate columnar bins oriented perpendicular to the cortical surface. Gray matter was further divided into fifteen equivolume depth bins based on local cortical thickness. Mean CO intensity was then computed within each columnar and depth bin, allowing us to characterize variation in metabolic architecture across both cortical surface position and depth.

A 3D histological volume was reconstructed from serial sections as previously described (2). In brief, brain tissue was segmented from the background using Illustrator (Adobe Systems Inc.) and converted to 8-bit grayscale images. To reduce computational demands while preserving resolution across cortical depth, images were downsampled to 768 × 1024 pixels for sagittal sections and 1024 × 768 pixels for coronal sections. Co-registration proceeded in two stages using MATLAB. First, a rigid-body alignment based on brain mask contours was applied across serial sections, using a middle slice as the reference. Second, gray matter was segmented from white matter using multilevel thresholding and used to refine alignment based on tissue contrast. These transformations were applied to the original CO-stained images, which were then stacked to form a 3D volume. To correct for inter-slice intensity fluctuations due to staining variability, we applied a section-wise intensity normalization procedure: we fit a quadratic curve to the mean section intensities and adjusted each slice to reduce deviations from the fitted trend, preserving gradual anatomical variation while minimizing technical artifacts.

### Creating and projecting (i) probabilistic face and place patches, (ii) eccentricity maps, and (iii) visual areas from probabilistic atlases

To define the face- and scene-selective regions in the IT cortex, we used a probabilistic group atlas of the middle lateral (ML) face patch and lateral place patch (LPP), generated from fMRI data in six monkeys for face and nine monkeys for place patches. These atlases were originally constructed in the National Institute of Mental Health macaque template (NMT) brain space (10, 33, 35). To align these maps with the histological data, we performed nonlinear registration between the NMT T1-weighted structural MRI and the reconstructed 3D histological volume using the JIP Analysis Toolkit (https://www.nitrc.org/projects/jip). Full details of this procedure are described in Oishi et al. (2024).

We used the same registration pipeline to project group-average probabilistic maps of retinotopic eccentricity and visual area boundaries, generated from six macaques (available at https://github.com/mikearcaro/BrainMaps/tree/main/Macaque/Retinotopy). To ensure that our eccentricity analyses focused on reliable and consistent responses, we applied spatial thresholds to include only voxels with a group-average F-statistic greater than 11 and a standard deviation in the eccentricity across macaques less than 3 degrees (Figure 2A). The histological sections included in this study span the occipitotemporal visual areas, including visual areas V1, V2, V3, V4, MT, PITd/v, OTS1/2. Face patch ML and place patch LPP largely overlap with PITd/v and OTS1/2, respectively (10, 11).

### CO profile comparisons between face and place patches

After projecting the face and place patches onto the histological sections, we quantified CO intensity within each region of interest (ROI). Because nonlinear warping can distort ROI boundaries and lead to incomplete coverage across cortical depth, we redefined each ROI directly in the histological volume by selecting columnar bins that intersected the projected ROI on the cortical surface and extending each bin through all 15 depth levels. This approach ensured complete sampling of cortical depth within each ROI across all sections.

Within each histological section, we averaged CO values across columnar bins within each depth bin, yielding a depth-resolved CO profile for both ML and LPP. To quantify overall metabolic activity per ROI per section, we then averaged these profiles across all 15 depth bins, resulting in a single section-level intensity measure. These section-level intensity values were compared between ML and LPP patches using unpaired, two-sample t-tests (Figure 1A). To assess whether differences were specific to cortical depth, we repeated this comparison for superficial (mean of bins 1–7) and deep (mean of bins 8–15) depth segments. All comparisons were FDR-corrected for multiple testing.

### Correlations between CO and eccentricity in face and place patches

To examine the relationship between CO intensity and retinotopic eccentricity within ML and LPP, we used each columnar bin within a patch as an individual data point. For each bin, we calculated mean CO intensity by averaging across all 15 depth bins. The corresponding eccentricity value was derived from the projected probabilistic eccentricity map. Because the fMRI-derived eccentricity map reflects a surface-based projection while CO data span cortical depth, we assigned each columnar bin a single eccentricity value based on its surface location. To combine data across monkeys, we first Z-scored the CO intensity and eccentricity values separately for each monkey (M1 and M2), then pooled the normalized data to calculate Pearson correlations between CO intensity and eccentricity across all columnar bins within each ROI.

### CO–Eccentricity Relationships Across Visual Cortex

To assess the relationship between CO intensity and eccentricity across the visual cortex, we computed the Pearson correlation coefficient across all columnar bins that overlapped with the group-average fMRI-defined eccentricity map. We fit both linear and second-order polynomial regressions between CO intensity and eccentricity for each monkey. To assess depth-specific effects, we repeated these analyses separately for the superficial (bins 1–7) and deep (bins 8–15) segments, averaging the Z-scored results across monkeys.

To identify visual areas that significantly deviated from the cortex-wide CO–eccentricity relationship, we computed standardized residuals from a robust linear regression (bisquare weighting; MATLAB’s nlinfit) of CO on eccentricity across all columnar bins. For each area (including ML and LPP), we calculated the mean of these residuals across its constituent columnar bins, separately for superficial and deep segments. With thousands of columnar bins per area, even tiny deviations from the expected relationship can achieve statistical significance despite representing negligible biological effects. Moreover, areas naturally vary in their intrinsic CO variability, making it difficult to distinguish genuine functional specialization from sampling noise or differences in measurement precision. To distinguish statistically detectable differences from biologically meaningful ones, we developed an eccentricity-matched null approach for each ROI. We binned each ROI’s eccentricity distribution into 0.5° intervals and sampled the exact same number of residual values from each bin across all other visual areas, repeated 5000 times to create stable null distributions. This preserved retinotopic structure while testing for area-specific metabolic patterns. We quantified practical significance using three complementary metrics:

1. within-ROI adjusted effect size standardized by the target ROI’s variance [d_adj = (μ_real - μ_null) /σ_real] with 95% bootstrap confidence intervals generated by resampling both distributions 1000 times with replacement,
2. distribution overlap computed as the area under the minimum of kernel density estimates [∫min(f_real(x), f_null(x))dx],
3. proportion of variance explained using eta-squared [η^2^ = SS_between /SS_total] where SS_between represents variance between real and null groups and SS_total represents total variance in the combined sample.

All metrics used identical null subsamples (matched to real ROI size) to ensure internal consistency. Areas were considered meaningfully different when showing low distributional overlap (<0.7), large adjusted effect sizes (|d_adj| > 1.0, indicating deviations exceeding the ROI’s intrinsic variability), and substantial variance explained (>5%), addressing the challenge that large sample sizes make traditional significance tests overly sensitive to trivial differences.

### Correction for tissue size effect

To rule out confounds related to potential staining variability due to the difference in tissue size across sagittal sections, we modeled and regressed out the effect of tissue size on CO intensity. We regressed CO intensity against tissue size of the sections across all columnar bins in the occipitotemporal cortex for each monkey. The resulting residuals, representing variance in CO independent of tissue size effect, were used in control analyses to confirm that the observed CO–functional relationships could not be explained by systematic staining artifacts. In this control analysis, we used raw CO intensity values without the white matter normalization described above (where we subtracted the difference between each section’s mean white matter intensity and individual gray matter pixel intensities) to avoid potential overcorrection that might artificially introduce or mask correlations between tissue size and CO staining patterns.

## Data and code availability

All CO histological sections, along with the aligned face and place patches and visual ROIs, are publicly available in the repository (https://osf.io/gjbmd/). This repository also contains the codes for generating the plots presented in the figures. Any additional information required to reanalyze the data reported in this paper is available from the lead contact upon request.

## Acknowledgments

This work was supported by a Whitehall Foundation grant (to M.J.A.) and by NIH grants EY16187 (to M.S.L.) and P30EY12196 (to M.S.L.). H.O. was supported by Toyobo Biotechnology Research Foundation long-term research grants and overseas research fellowships of JSPS. K.S.W. was supported by the Brain and Behavior Research Foundation (NARSAD 30738) and the National Science Foundation (NSF CAREER 204225). The funders had no role in the study design, data collection and interpretation, or the decision to submit the work for publication.

