## Supplementary material for "Metabolic organization of macaque visual cortex reflects retinotopic eccentricity and category selectivity": Figure S1 and Table S1

1) Dept. Psychology, UC Berkeley, Berkeley, CA, USA, 2) Div. of Sensory and Cognitive Brain Mapping, Dept. System Neuroscience, National Institute for Physiological Sciences, Okazaki, Aichi 444-8585, Japan, 3) Dept. Neurobiology, Harvard Medical School, Boston, MA, USA, 4) Dept. Neuroscience, UC Berkeley, Berkeley, CA, USA, 5) Helen Wills Neuroscience Institute, UC Berkeley, Berkeley, CA, USA, 6) Dept. Psychology, University of Pennsylvania, Philadelphia, PA, USA

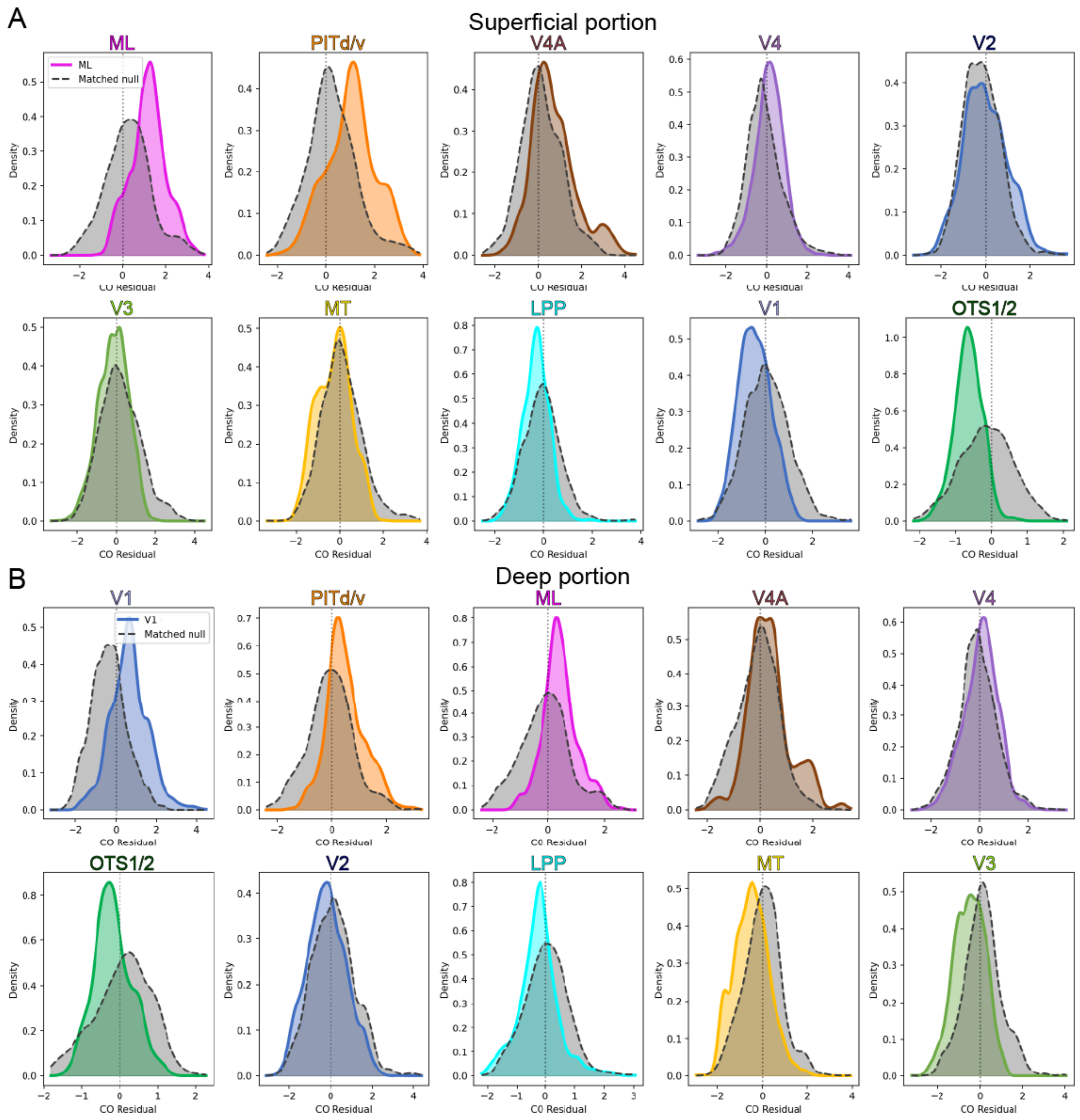


**Fig. S1 CO residual distributions for each visual area in superficial (A) and deep portions (B).** Each panel displays the overlap between CO residual distribution for each visual area (colored by ROI) and the eccentricity-matched null distribution (gray). Panels are ordered by mean residual magnitude of the areas (highest to lowest). The distributions are identical to those shown in Figure 4B.

| Depth | ROI | Adjusted effect size | CI_Low | CI_High | Overlap | Variance explained |
|---|---|---|---|---|---|---|
| Superficial portion | ML | 1.100 | 1.026 | 1.176 | 57.6% | 19.1% |
| | PITd/v | 0.782 | 0.716 | 0.851 | 62.0% | 13.8% |
| | V4A | 0.542 | 0.497 | 0.588 | 79.3% | 7.5% |
| | V4 | 0.280 | 0.247 | 0.312 | 77.3% | 1.6% |
| | V2 | 0.156 | 0.124 | 0.189 | 89.6% | 0.7% |
| | V3 | -0.493 | -0.527 | -0.460 | 83.3% | 4.1% |
| | MT | -0.537 | -0.601 | -0.474 | 82.8% | 5.4% |
| | LPP | -0.452 | -0.539 | -0.366 | 79.3% | 3.4% |
| | V1 | -0.698 | -0.738 | -0.658 | 74.9% | 8.0% |
| | OTS1/2 | -1.246 | -1.402 | -1.098 | 57.6% | 15.2% |
| Deep portion | V1 | 1.070 | 1.035 | 1.105 | 54.5% | 23.3% |
| | PITd/v | 0.816 | 0.746 | 0.886 | 67.1% | 12.5% |
| | ML | 0.893 | 0.819 | 0.970 | 66.4% | 12.0% |
| | V4A | 0.457 | 0.409 | 0.506 | 82.5% | 5.1% |
| | V4 | 0.219 | 0.188 | 0.250 | 85.7% | 1.1% |
| | OTS1/2 | -0.461 | -0.585 | -0.343 | 71.1% | 3.4% |
| | V2 | -0.315 | -0.353 | -0.277 | 86.9% | 2.2% |
| | LPP | -0.376 | -0.452 | -0.303 | 74.8% | 3.3% |
| | MT | -0.656 | -0.718 | -0.595 | 71.7% | 9.1% |
| | V3 | -0.853 | -0.886 | -0.820 | 70.8% | 13.0% |

**Table. S1 Statistical difference between CO residual and eccentricity-matched null distributions for each visual area in Figure 4B and S1.** Adjusted effect size represents the standardized d-prime and CI_Low and CI_High represent the 95% bootstrap confidence intervals. Overlap represents the proportion of the distribution overlap, computed as the area under the minimum of kernel density estimates. Variance explained represents the proportion of eta-squared. See Materials and Methods for the details of the indices. Data are ordered by mean residual magnitude of the visual areas (highest to lowest).
